# Evolutionary tradeoffs and the structure of allelic polymorphisms

**DOI:** 10.1101/244210

**Authors:** Hila Sheftel, Pablo Szekely, Avi Mayo, Guy Sella, Uri Alon

## Abstract

Populations of organisms show prevalent genetic differences called polymorphisms. Understanding the effects of polymorphisms is of central importance in biology and medicine. Here, we ask which polymorphisms occur at high frequency when organisms evolve under tradeoffs between multiple tasks. Multiple tasks present a problem, because it is not possible to be optimal at all tasks simultaneously and hence compromises are necessary. Recent work indicates that tradeoffs lead to a simple geometry of phenotypes in the space of traits: phenotypes fall on the Pareto front, which is shaped as a polytope: a line, triangle, tetrahedron etc. The vertices of these polytopes are the optimal phenotypes for a single task. Up to now, work on this Pareto approach has not considered its genetic underpinnings. Here, we address this by asking how the polymorphism structure of a population is affected by evolution under tradeoffs. We simulate a multi-task selection scenario, in which the population evolves to the Pareto front: the line segment between two archetypes or the triangle between three archetypes. We find that polymorphisms that become prevalent in the population have pleiotropic phenotypic effects that align with the Pareto front. Similarly, epistatic effects between prevalent polymorphisms are parallel to the front. Alignment with the front occurs also for asexual mating. Alignment is reduced when drift or linkage is strong, and is replaced by a more complex structure in which many perpendicular allele effects cancel out. Aligned polymorphism structure allows mating to produce offspring that stand a good chance of being optimal multi-taskers in at least one of the locales available to the species.

## Media Summary

Populations of organisms show many genetic differences called polymorphisms. These polymorphisms often cause variation in biological traits between individuals, and thus understanding them is important for biology and medicine. Here, we ask which polymorphisms occur when organisms evolve under tradeoffs between multiple tasks. Multiple tasks present the organism with a problem, because it is not possible to be optimal at all tasks at once and hence compromises are necessary. We use theory and simulation to find that polymorphisms under multi-task natural selection have special structure that ‘learns’ the range of optimal compromises between the tasks. This special structure allows mating to produce offspring that stand a good chance of being optimal multi-taskers.

## Introduction

Organisms show prevalent genetic differences called allelic polymorphisms. Allelic polymorphisms, together with environmental and epigenetic effects, cause much of the variation in traits between individuals. Each polymorphism typically affects multiple phenotypic traits at once (pleiotropy), and each trait is usually affected by many different polymorphisms [1,2].

Understanding the origin and distributions of polymorphisms is of central importance in evolutionary and quantitative genetics. Most theoretical approaches to the evolution of polymorphisms employ the classic picture of the fitness landscape: a genotype determines phenotype, and the phenotype determines fitness [3] (Fig 1A). Natural selection leads to phenotypes that maximize fitness, provided that there is sufficient time, genetic variation, and that populations and selection pressures are large enough to overcome genetic drift [4]. In this approach, the shape of the fitness landscape, in particular the slopes near the maximum, influences the distribution of phenotypes and genotypes in the population. For example, simulations show that a fitness ridge can lead to mutation-selection balance in which traits vary along the ridge [5,6].

**Figure 1.**
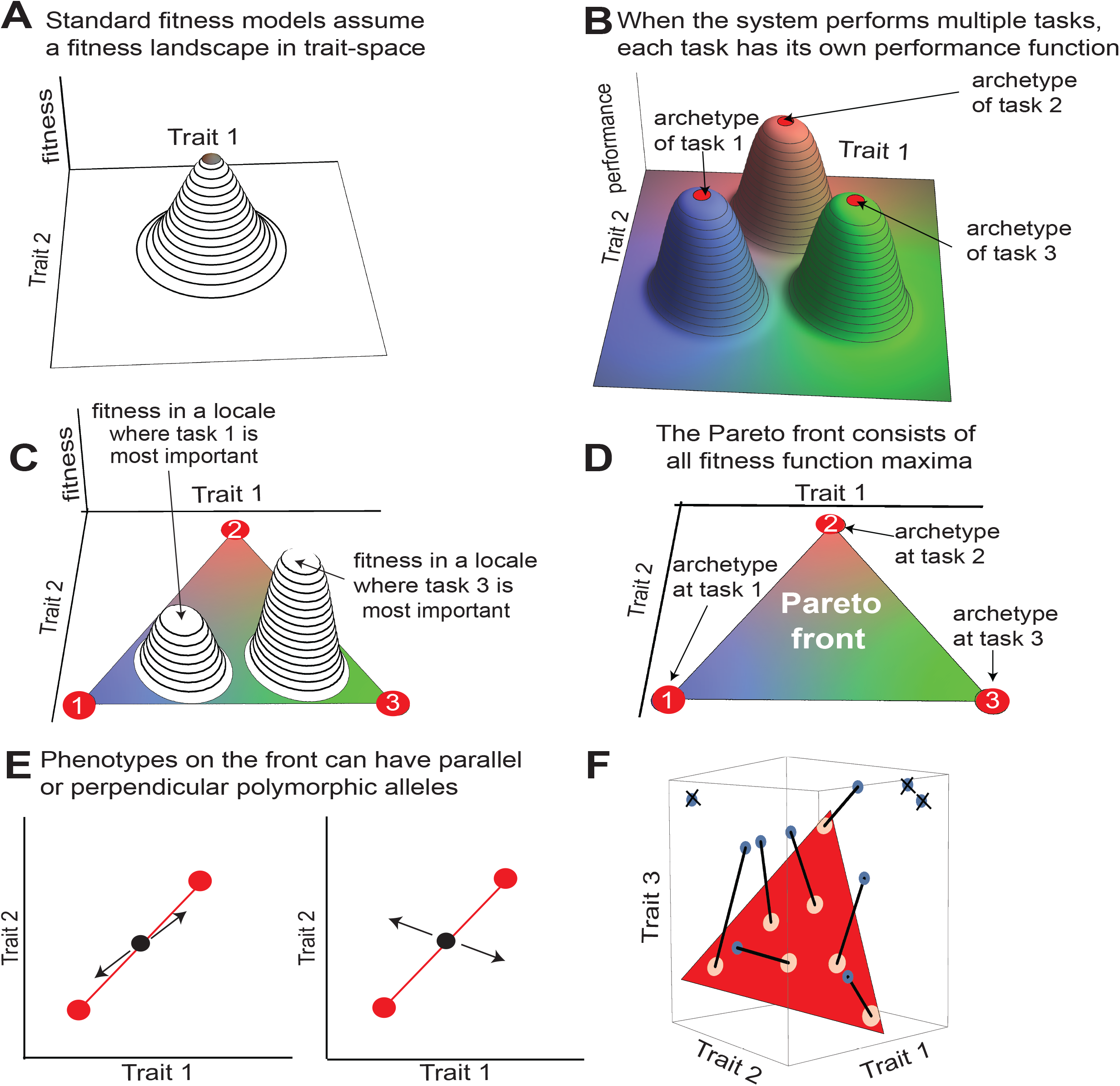
Tradeoff between tasks leads to phenotypes arranged along polygons in trait space. **(A)** Standard approaches assume a fitness landscape in the space of phenotypic traits. Selection tends to favor phenotypes near the maximum. **(B)** When the system needs to perform several tasks, each task has a performance function in trait space, whose maximum is called the archetype. Each colored hill represent a performance function 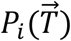 at a single task i, where 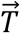 is the vector of traits, and fitness at locale q is an increasing function of all performance functions 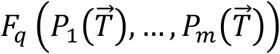 **(C)** In an environment where task 1 is more important, fitness is maximized at a phenotype close to archetype 1; In an environment where task 3 is more important, fitness is maximized at a phenotype closer to archetype 3. Fitness maxima in the two environments can be different. **(D)** Fitness maxima in all possible environments in which fitness is an increasing function of performance in the three tasks lie in the full triangle whose vertices are the 3 archetypes-known as the Pareto front-when the conditions of Shoval et. al [9] are fulfilled. **(E)** Possible allelic polymorphism structures that keep a phenotype on the front: either mutation effects are parallel to the front (left), or pairs of mutations whose components perpendicular to the front cancel out (right). **(F)** Schematic of the selection process described in the Results section, for the case of three tasks that define a triangle whose vertices are the three archetypes. Locales and individuals best at the locales are sequentially removed into a survivor list, and remaining individuals are removed. Survivors make up the next generation. Here we consider 7 locales (N=7); phenotypes that maximize fitness in these locales are connected to the relevant locale using a black line.

Here, we consider an extension of the fitness landscape picture for cases in which fitness derives from an organism’s performance at *multiple tasks*. The need to perform multiple tasks introduces an inherent tradeoff, because in most cases no single phenotype can be optimal at all tasks. For example, a bird may need to both eat seeds and pollen. However, cracking seeds requires a beak shaped like a plier whereas pollen requires a pincer-like beak [7].

In such cases, the ‘genotype → phenotype → fitness’ scheme should be modified to include performance at different tasks as an intervening step between phenotype and fitness [8]:genotype → phenotype → performance at task 1,2,…➔ fitness). The genotype determines phenotype, which is defined by a set of traits 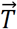 such as morphological parameters (e.g. beak width and length), gene expression levels or enzyme activities. The phenotype, in turn, determines the performance at different tasks: The performance at task *i* is given by the performance function 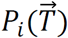 (Fig 1B). Fitness is an increasing function of the performances, *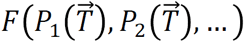*. The fitness in a given locale is given by some (potentially nonlinear) weighting of the performances in the different tasks (Fig 1C). Therefore, each locale has its own fitness function F that combines the performances in the tasks.

This picture of tradeoffs between tasks has been shown to result in a simple geometry of phenotypes in the space of traits (trait space) [9]. Under quite general assumptions, the optimal phenotypes in all possible environments fall within polytopes in trait-space (polytopes are the generalization of polygons to any dimension, such as lines, triangles, tetrahedra, etc.). The vertices of the polytopes are the phenotypes that are optimal for a single task. These vertices are called *archetypes*, and the polytopes they define are the *Pareto fronts*.

Each point on the Pareto front corresponds to a phenotype that maximizes fitness in a locale that requires a specific degree of specialization in each of the tasks (Fig 1C). For example, a trade-off between two tasks leads to phenotypes along a line segment, with the two archetypes at either end; generalists lie in the middle. A trade-off between three tasks leads to a full triangle (Fig 1D), four tasks lead to a tetrahedron and so on. Such straight edged polytopes occur in coordinate systems in which performance declines with a distance metric [10]. If trait axes are nonlinearly transformed (e.g. trait->trait^2^), the polygons have curved edges, but still preserve the distinct vertices at the archetypes optimal for each task.

Evidence for such lines, triangles and tetrahedra was found in morphological datasets [9,11,12]; animal life-history traits [13]; locomotive behavior [14,15] and gene expression data [9,16–18]. The polytopes offer a way to deduce the tasks from the data, by observing the special features of organisms near each archetype. In the studies cited above, the archetypes found by fitting the data to polytopes [16] revealed clues about the tasks at play in each case. For example, primary tasks for *E. coli* gene expression are growth and survival [9]. For this reason, this theoretical approach is called Pareto task inference (ParTI).

Up to now, work on the ParTI approach has not considered its genetic underpinnings. Here, we address this by asking how the allelic polymorphism structure of a population is affected by evolution under tradeoffs between multiple tasks. We simulate a multi-task selection scenario, in which the population evolves to the Pareto front: the line segment between the two archetypes or the triangle between three archetypes. We find that the polymorphisms that become prevalent in the population have phenotypic effects that align with the Pareto front. Epistatic effects between prevalent polymorphisms are also parallel to the front. This polymorphism structure allows rapid evolution to new environments that require the same tasks at different weightings. It also provides a mechanism that allows mating to produce offspring that stand a good chance of being optimal multi-taskers. Alignment to the front is reduced when genetic drift is strong, and is replaced by a more complex structure in which perpendicular effects of alleles collectively cancel out.

## Materials and Methods

We assume that performance at each task decays with distance *r* to the archetype [9,19]. Performance functions were 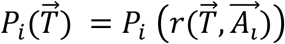, where distance *r* from the archetype 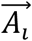 is an inner-product 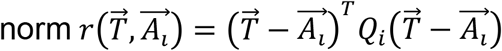 with a positive-definite matrix Q_i_ (Euclidean distance is obtained for Q_i_ = *I*). Unless otherwise noted, all simulations used Q_i_ = *I*. We chose population sizes N for which simulation duration for 10N generations was feasible, namely up to N=2000.

## Results

### A selection scheme for multi-task evolution

We consider a setting where fitness is determined by the performance at *L* different tasks. Phenotypes are described as vectors 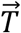 in a trait space, where each axis corresponds to a quantitative trait. The performance functions for the *L* tasks are 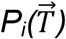 for *i=1.L*. The maxima of these performance functions are the archetypes, 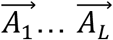, the phenotypes optimal at single tasks, and performance at each task decays with distance to the archetype [9,19] (Methods).

To address evolution under a tradeoff between the tasks, we developed a selection process as follows: we consider N discrete locales, such as territories or nesting sites, each of which can be occupied by a single individual (results apply also when locales can be occupied by several individuals, Electronic Supplementary Material, Section 1). The locales differ in environmental factors, and therefore the different tasks are more or less important for overall fitness at the locale. This heterogeneity can result from geographic clines, patchy environments, variation in other species, and so on. Hence, each locale has its own fitness function (individual selection surface) with locale-specific weights for the different performance functions: 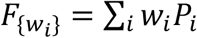 where the weights *w_i_* are positive (similar results are found for fitness functions that are non-linear in the performances, Electronic Supplementary Material, Section 2). In the simulations shown below, we sample w_i_ uniformly with Σ*w_i_* = 1. Other distributions for w_i_ yield the same qualitative results (Electronic Supplementary Material, Section 3).

The weights *w_i_* correspond to the fitness contribution of each task in the locale. With this mathematical description, the phenotype that maximizes fitness in each locale can be shown to fall within trait space in the polytope whose vertices are the archetypes [9], namely the Pareto front: a line for two tasks, a triangle for three tasks and so on.

Each round of the simulation begins with *N* diploid individuals, each in a different locale. We randomly choose *kN* pairs (*k* > 1) and mate each pair with recombination to generate one offspring per pair (simulations where mating probability depends on fitness yield the same qualitative result, Electronic Supplementary Material, Section 4). Offspring gain new mutations with probability *μ*.

The *kN* offspring compete for the locales as follows. We choose a locale at random, select the individual with highest fitness according to the fitness function for that locale. We put that locale and individual aside in a ‘survivor list’, and repeat the process with remaining individuals and locales until all sites are filled. The remaining individuals are removed, and the *N* survivors make up the next generation (Fig 1F). Selecting individuals stochastically according to their fitness yields the same quantitative results, Electronic Supplementary Material, Section 5). As the parameter *k* increases, the competition for the sites intensifies and drift effects become smaller.

This selection scheme acts on individuals represented by a genome composed of a set of mutant alleles. Each mutation moves the phenotype in trait-space. Mutation effect is a randomly oriented vector whose length is drawn from an exponential distribution with mean q=1, which is 1% of the distance in trait space between the archetypes. Mutation effects are additive (non-additive epistasis is addressed below). The number of new mutations introduced in an individual per generation is Poisson distributed with mean μ. For recombination we assume an infinite site model [20,21] and free recombination [22] (for more details see Electronic Supplementary Material, Section 6).

We also simulated variants of this model as follows: (i) linkage, in which recombination was done by swapping chromosomes at a single recombination spot (ii) No mating: asexual populations with a single chromosome and no recombination.

### Phenotypes in the population become spread along the Pareto front

We begin with simulations with two tasks, which define a Pareto front in the shape of a line segment between the two archetypes. The initial condition is a cloud of phenotypes that are off the Pareto front (Fig 2A). For a wide range of model parameters (*Nμ* =10-50, N=100-2000, k=1.1−5), we find that within a few tens of generations the population spreads along the Pareto front. The distribution around the line becomes narrower as generations pass (Fig 2B-E).

**Figure 2.**
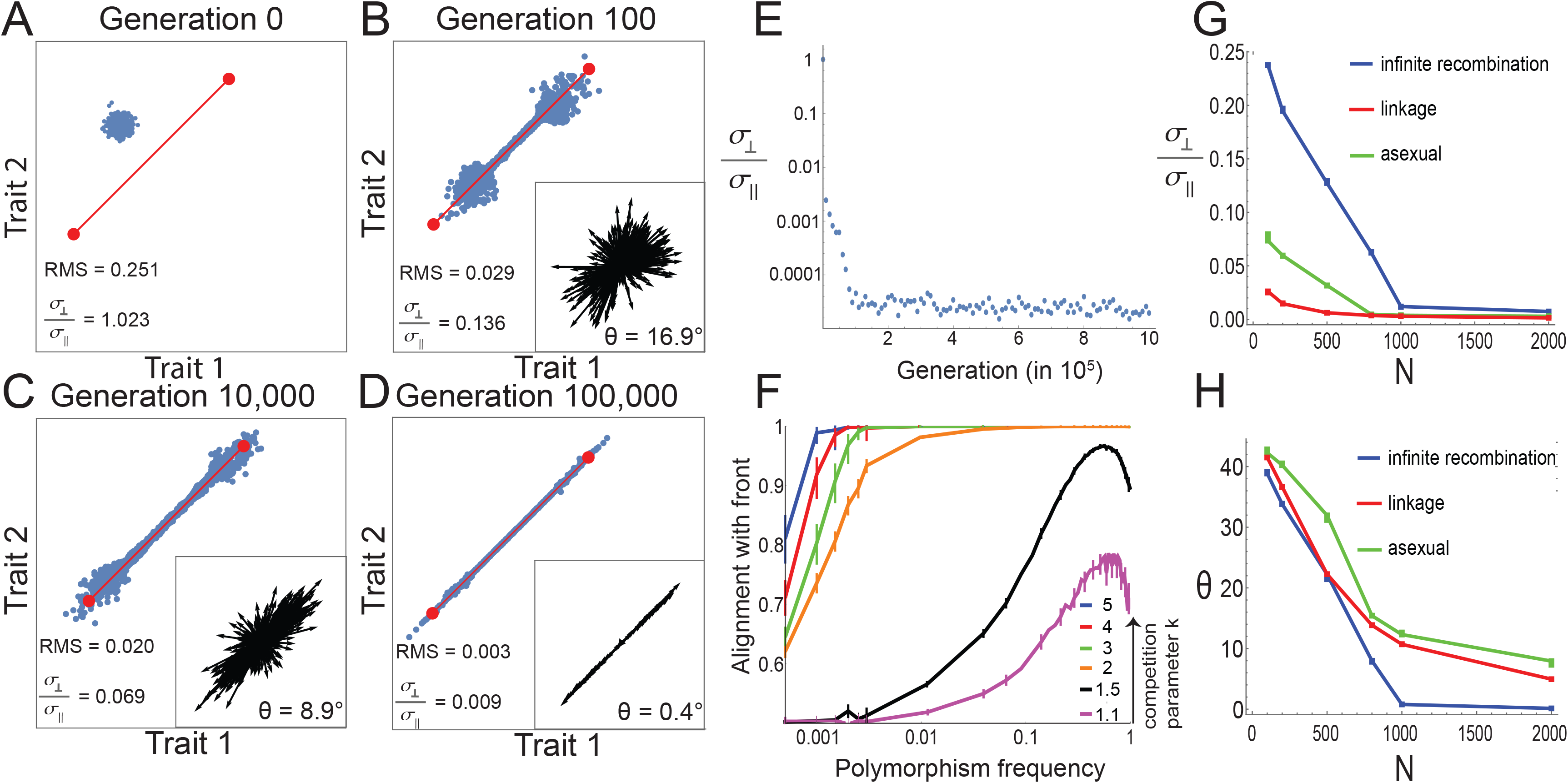
Phenotypes converge to the Pareto front by means of polymorphisms whose effects align with the front. **(A-D)** Snapshot of the phenotypes at different generations in a simulation, for the case of two tasks. The Pareto front is shown in red. Insets: The phenotypic effects of all allelic polymorphisms present in at least 1% of genomes increasingly align with the Pareto front. Each arrow represents the phenotypic effect of one polymorphism (magnified for illustration purposes). At generation zero, no mutation is present at >1% of the genomes. Simulation parameters: *N* =1000, *μ*=0.05, *k* = 2, infinite recombination. Median angle *θ* of polymorphisms relative to the front was 17°, 9°, 0.4°, for B-D, respectively. The ratio between perpendicular and parallel standard deviation of phenotypes is mentioned in each panel A-D, along with the phenotypic RMS distance from the front, normalized by the distance between archetypes. Axes are traits 1 and 2. **(E)** The ratio of the perpendicular and parallel standard deviation of phenotypes with respect to the front is 1 for the initial population at generation 0, decreases with generations, and begins to plateau after ~100,000 generations. **(F)** Polymorphisms that are aligned with the front tend to increase in frequency (log-linear scale) and with competition k. Simulation parameters are as above except k that varies as indicated. Error bars represent 95% confidence intervals from bootstrapping. Alignment is defined as 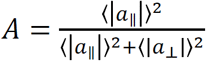, where 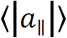 and 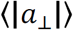 are the mean parallel and perpendicular component of the mutation effect vectors in each bin of mutation frequency. A = 1 and 0.5 occur when mutations are completely aligned or randomly oriented, respectively. Note that data is for frequencies <1, at frequency=1 the mutation becomes fixed in the population. **(G-H)** Phenotypic standard deviation ratio 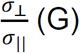 (G) and median angle *θ* (H) to the front, as a function of population size N, for three different recombination schemes: free recombination, linkage, and asexual mating. In all simulations presented, *μN*=50, and competition parameter k =2. Error bars represent 95% confidence intervals from bootstrapping.

Regardless of parameters, phenotypic standard deviation parallel to the front *σ*_||_ is much larger than perpendicular standard deviation *σ*_⊥_. For example, for N=1000 and k=2, we find after 100N generations *σ*_⊥_/*σ*_||_ = 0.012 ± 0.0003. Similar results are found also when varying the shapes of the performance function contours *(Q_i_* ≠ *I*), and for linkage and asexual mating, as described in Electronic Supplementary Material, Section 7 and in Fig 2G.

### Prevalent polymorphisms have effects parallel to the Pareto front

We next asked about the allelic polymorphism structure of the evolved population. In principle, there are two possible structures that give rise to phenotypes along the front: either mutation effects are parallel to the front, or there exists pairs of mutations or higher-order combinations whose components perpendicular to the front cancel out (Fig 1E).

We measured alignment with the front using the angle *θ* of each allele effect with the front. We find that while newly arising mutations are isotropic (median angle=45°), polymorphisms that persist in the population (>1% of the population) are closely aligned with the Pareto front (Fig 2B-D insets). For example, the median angle of these polymorphisms is 0.80° ± 0.03° after 100,000 generations, for typical simulation parameters (N=1000, k=2, *μ*=0.05).The more aligned a polymorphism is with the front, the more frequent it is in the population (Fig 2F, Electronic Supplementary Material, Sections 8–9). We next asked how genetic drift affects the alignment. We find that the smaller the population (the larger the drift), the weaker the alignment (Fig 2H, Electronic Supplementary Material, Section 8). Similarly, alignment is weaker when the competition parameter k is small, again a situation with larger drift (Fig 2F). Drift effects seem to become important at population sizes smaller than a few hundreds.

We also asked about the effects of recombination on alignment. Alignment is reduced at a given population size when free recombination is replaced with single-site recombination (linkage) or in the case of asexual mating (no recombination) (Fig 2H). At small populations sizes alignment is poor (>20° at N=500, for both asexual mating and linkage) but the phenotypes are still are close to the front (*σ*_↖_/*σ*_||_ < 0.05), indicating that the allelic polymorphisms have a complex structure in which perpendicular effects cancel out. Nevertheless, simulations with linkage or asexual mating still showed good alignment (angle<10°) at large population sizes.

We conclude that regardless of linkage or asexual mating, at large enough population sizes common allelic polymorphisms align with the front in these simulations, whereas at small population sizes alignment is weaker and instead polymorphism structure is more complex with cancelling perpendicular effects.

### Phenotypic effects of polymorphisms align with moderately-curved Pareto fronts

We also tested curved Pareto fronts which occur when the performance functions have eccentric contours *(Q_i_* ≠ *I*) that point at an angle with respect to each other [10], as shown in Electronic Supplementary Material, Section 10, Fig S13. When the front curvature is mild, polymorphisms are still aligned with the local front (for example, when k=2 and curvature is as shown in Fig S11C, the median polymorphism angle is 6.1° ± 0.1°); this alignment is reduced at high front curvatures and high competition (for example, when k=5 and curvature is higher as shown in Fig S12B, the median angle is 24.1° ± 0.2°) (Fig S11–13, Electronic Supplementary Material, Section 10).

### Allelic polymorphism effects align with a triangular Pareto front in the case of three tasks

We asked whether these conclusions apply also to higher numbers of tasks. We simulated evolution under three tasks, which results in a Pareto front shaped as a triangle whose vertices are the three archetypes. We used a three-dimensional trait space, with the triangle oriented at angle with respect to all trait axes. We find that, again, populations rapidly converge to the triangle (Fig 3A-C). Prevalent polymorphisms align with the plane of the triangle (median angle ~2°, Fig 3D-F). This effect is seen also when mating probability depends on fitness (Electronic Supplementary Material, Section 11).

**Figure 3.**
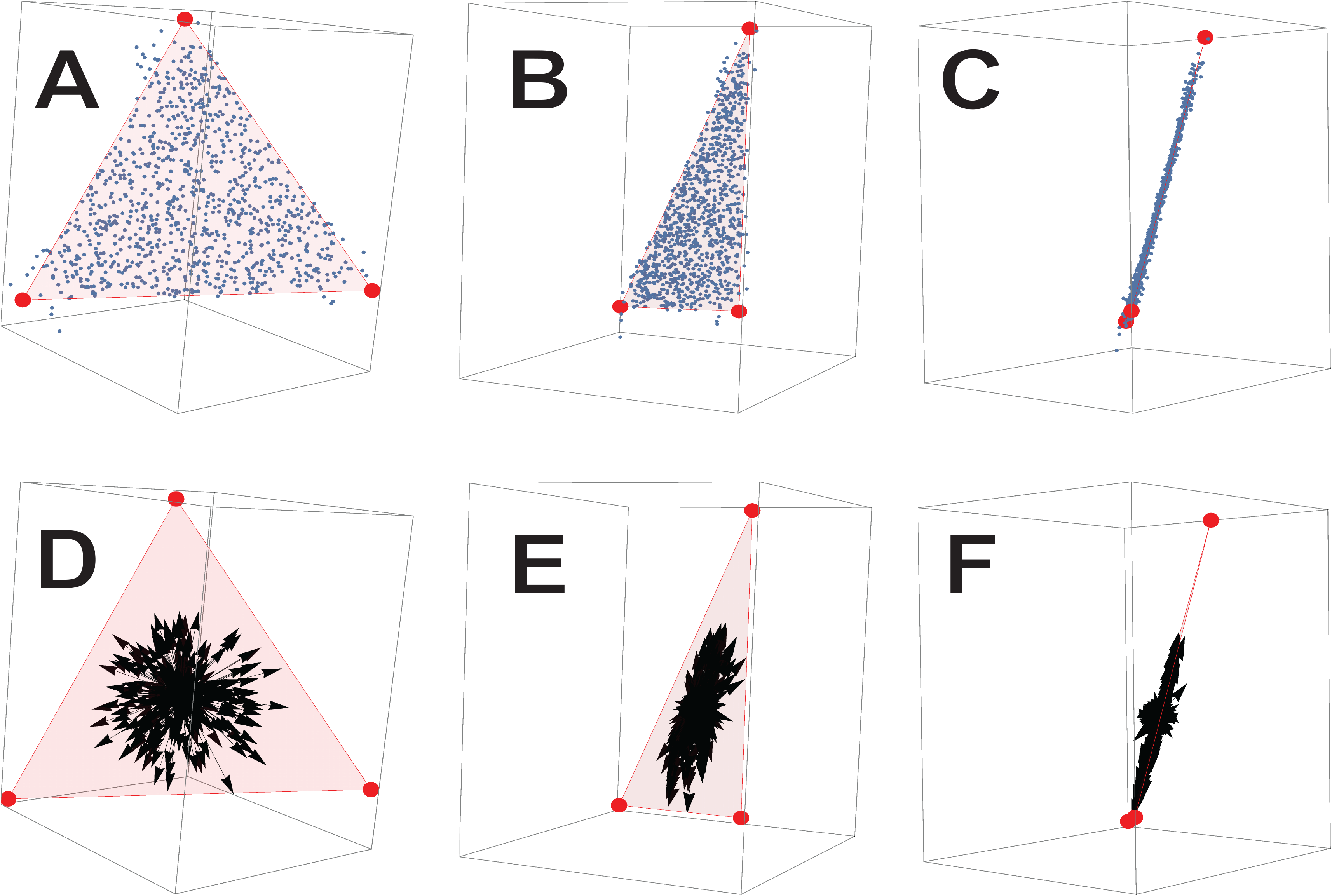
For three tasks, polymorphisms align with the plane of the triangular Pareto front. **(A-C)** Phenotypes (blue dots) lie close to the Pareto (red triangle) front after 100,000 generations. Panels show the data from three different views. Simulation parameters are N = 1000, *μ*=0 05, k=5, infinite-recombination. **(D-F)** Effects of allelic polymorphisms present in at least 1% of the genomes (black arrows) align with the Pareto front (effects magnified by 10-fold for visualization). Each arrow represents the phenotypic effect of one polymorphism. Panels show the data from three different views.

### Epistatic effects between the two copies of the same allele align with the Pareto front

We next consider the effects of tradeoffs on epistasis. We begin with the non-additive interaction between two alleles of the same gene [23–25], which is related to the phenomenon of dominance. We modelled this epistasis by assigning to each mutation *i* a randomly oriented vector 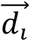, such that the heterozygote mutant effect is 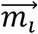 and the homozygote is 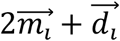. We find that as before, the main effects of prevalent polymorphisms 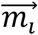 align with the front. Importantly, the epistatic effects 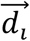 of common polymorphisms also align with the front (Electronic Supplementary Material, Section 12). Those with epistatic effects off of the front are selected against. Thus epistasis, not only main effect, tends to align with the front, provided that drift is not too large.

### Epistasis in a molecular mechanism that generates triangular fronts

In addition to epistasis between the alleles of the same gene, we studied epistasis between different genes. In this case, the effects of each polymorphism depend on the genetic background. To study this, we focus on epistasis due to nonlinear interactions within a molecular mechanism inspired by gene expression [9]. This molecular mechanism is of interest because it occurs in bacteria, and naturally provides scope for a triangular Pareto front.

In the model, genes are regulated by three regulators X_i_ that compete over a limiting factor R. For example, in bacteria, three sigma factors compete over RNA polymerase [26,27]) (Fig 4A). The traits are the expression of different genes. The genes are regulated by the three sigma factors bound to RNA polymerase which bind to sites in the promoter of the gene. The effect of regulator i on gene j expression is *ω_ij_*. Thus, expression of gene j is 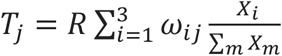, (Electronic Supplementary Material, Section 13). The competition between X_i_ for binding to R leads to the nonlinear term 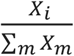 which causes epistasis between allelic variants.

**Figure 4.**
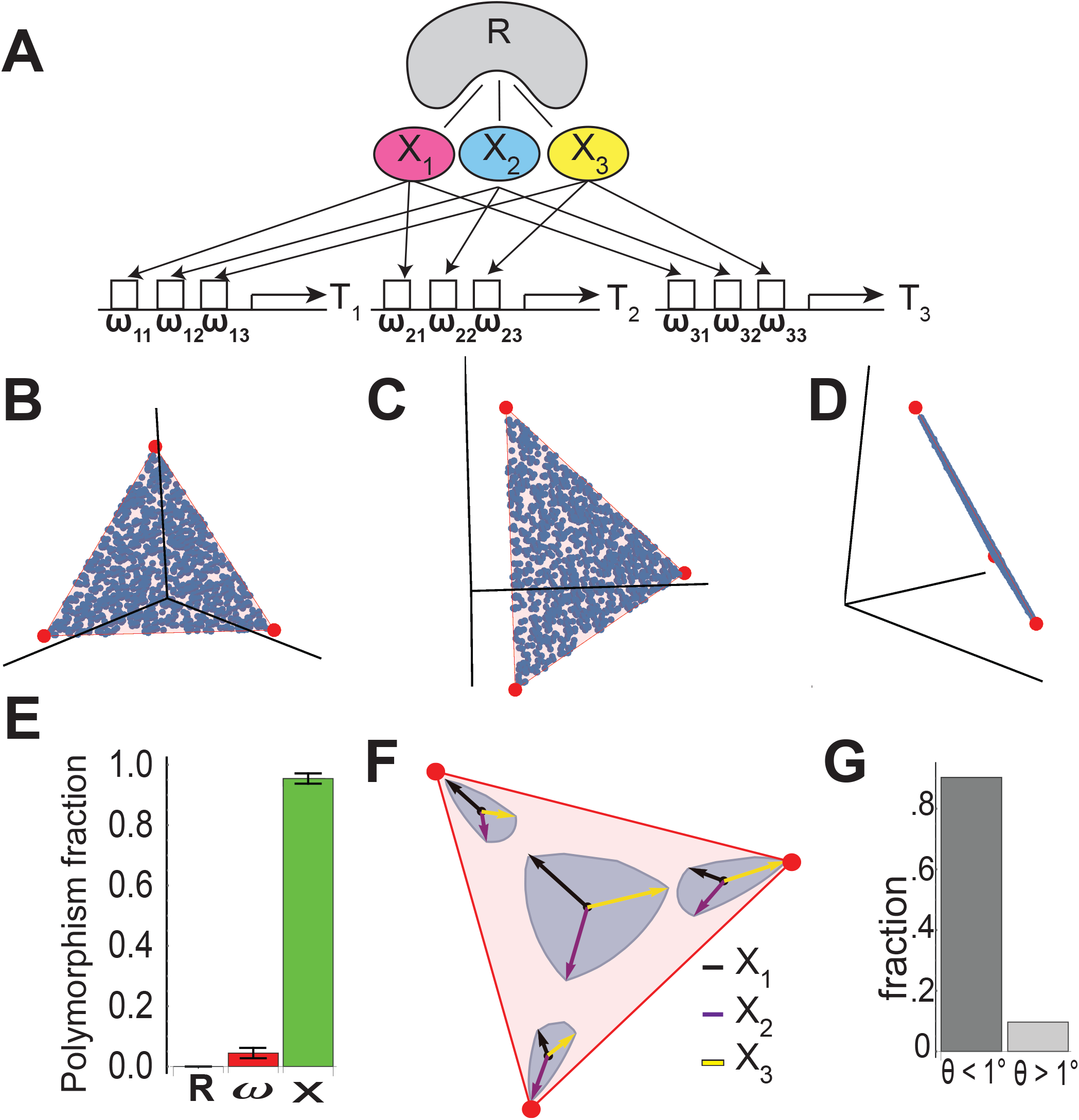
Epistasis due to a molecular mechanism evolves to keep phenotypes within the triangular front. **(A)** A mechanism based on bacterial transcription considers genes whose promoters are regulated by 3 regulators (sigma factors) X_i_ that compete over a limiting factor R (RNA polymerase). The effect of X_i_ on the promoter of gene j is ω_ij_. **(B-D)** Phenotypes (blue dots) reside on the Pareto (red triangle) front after 100,000 generations with this mechanism. Panels show the data from three different views. Simulation parameters (N = 1000, *μ*=0.05, k=5, equal maximal fitness in all locales. Each mutation changed one of the parameters X_i_, ω_ij_ or R by multiplying the parameter by number drawn from a lognormal distribution LN(1,1). **(E)** Fraction of prevalent polymorphisms (>1% of genomes) in each of the parameters R, ω_ij_ (summed over all ω’s) and X_i_ (summed over X_1_, X_2_, X_3_). Median and median deviation over 100 different simulations is shown at 100,000 generations. **(F)** Effect of mutations that change X_i_ by 10%. Arrows are 10% changes in X_1_ (black arrow), X_2_ (purple arrow), or X_3_ (yellow arrow), and the blue contour is the boundary of the effects of simultaneously changing X_1_, X_2_ and X_3_ so that the total change is 10%. Mutation effects are magnified by 12 for illustration. Mutations show epistasis, by having different effects at different genetic backgrounds that correspond to four different phenotypes on the front. **(G)** Fraction of angles with respect to the plane of the triangle that are smaller (left) or bigger (right) than 1°, for all polymorphisms present at >1% of the genomes. Each polymorphism was evaluated in all of the genomes in which it appears, because the angle can vary due to epistasis. Results in (B-D) are based on simulation with parameters N = 1000, *μ*=0.05, k=5, infinite-recombination. Each mutation changed one of the parameters X_i_, ω_ij_ or R by multiplying the parameter by number drawn from a lognormal distribution LN(1,1).

We simulated evolution in this model for the case of three tasks (in bacteria such as *E. coli*, tasks can be growth, survival and motility regulated by the σ-factors σ_70_, σ_S_ and σ_A_ [26,27]). Each mutation varies one of the biochemical parameters: X_i_, *ω_ij_* or R. Phenotypes evolve to the triangular front (Fig. 4B-D).

Importantly, we find that almost all of the prevalent polymorphisms affect the levels of the regulators X_i_; there are almost no prevalent polymorphisms that affect the levels of the other biochemical parameters, the promoter strengths *ω_ij_* or RNA polymerase levels R (Fig 4E), once the mutations that set *ω_ij_* become fixed (at around 1000 generations, Electronic Supplementary Material, Section 14).

Thus, evolution first encodes the coordinates of the three archetypes in the promoter sequences that set the weights *ω_ij_*. The position on the triangle is then given by the relative values of the regulators X_i_. For example, phenotypes at the vertices of the triangle have one regulator X_i_ high and the rest very low.

The polymorphisms in X_i_ have phenotypic effects that depend on the genetic background (Fig 4F) due to the epistasis. For genetic backgrounds near the center of the front, polymorphisms in X_i_ move the phenotype in all directions along the plane of the triangle; near the vertices, they move the phenotype in a tapered way that fits into the corner of the triangle, preventing phenotypes from leaving the triangle. None of these polymorphisms has a sizable component that moves off of the plane of the triangle (Fig 4G). In contrast, the mutations that are actively selected against and hence do not reach high prevalence - those in the parameters *ω_ij_* (promoter mutations) and R (RNA polymerase) - have effects that move the phenotype off of the plane the triangle. For example, increasing a weight *ω_ij_* moves expression of gene i up, without affecting the other genes, a move which is off of the plane of the triangle. Altering the level of the limiting substrate R changes expression of all genes at once, a direction that is also off of the plane of the triangle. Expression levels of R and of genes will presumably be determined by the benefit and cost of production and maintenance. Such cost is included in the performance functions - increasing expression of genes carries a cost that, at high enough expression levels, reduces performance [28,29]. This demonstrates how a regulatory mechanism focuses the selection of mutations to certain components (regulator activity) and not to others (promoters, RNA polymerase). The common polymorphisms have epistatic interactions that keep them within a sharp triangle of phenotypes [30]. The same idea can be generalized to any number of tasks by using more regulators.

### Polymorphism structure under tradeoffs allows rapid selection along the Pareto front

The polymorphism structure found above has implications for artificial selection towards desired traits. To explore this, we exposed populations that have evolved for 100K generations under two or three tasks (linear or triangular Pareto front) to selection towards a given phenotypic target (Electronic Supplementary Material, Section 15). We compared selection to a target aligned with the original front to selection to a target that is not aligned with the front. Each generation, we selected the top fraction *p* of the population closest to the target.

We find that evolution to a target aligned with the original front is much faster and results in phenotypes that move farther from the original phenotypes than selection perpendicular to the original front (Fig 5A-B and Electronic Supplementary Material, Section 15, Fig S17B). For example, for typical 3-task simulation parameters (N=1000,k=5,*μ*=0.05), and artificial selection parameter p in the range 0.1–0.9, response to selection aligned with the original front is 9-fold to 17-fold larger than the response to selection perpendicular to it.

**Figure 5.**
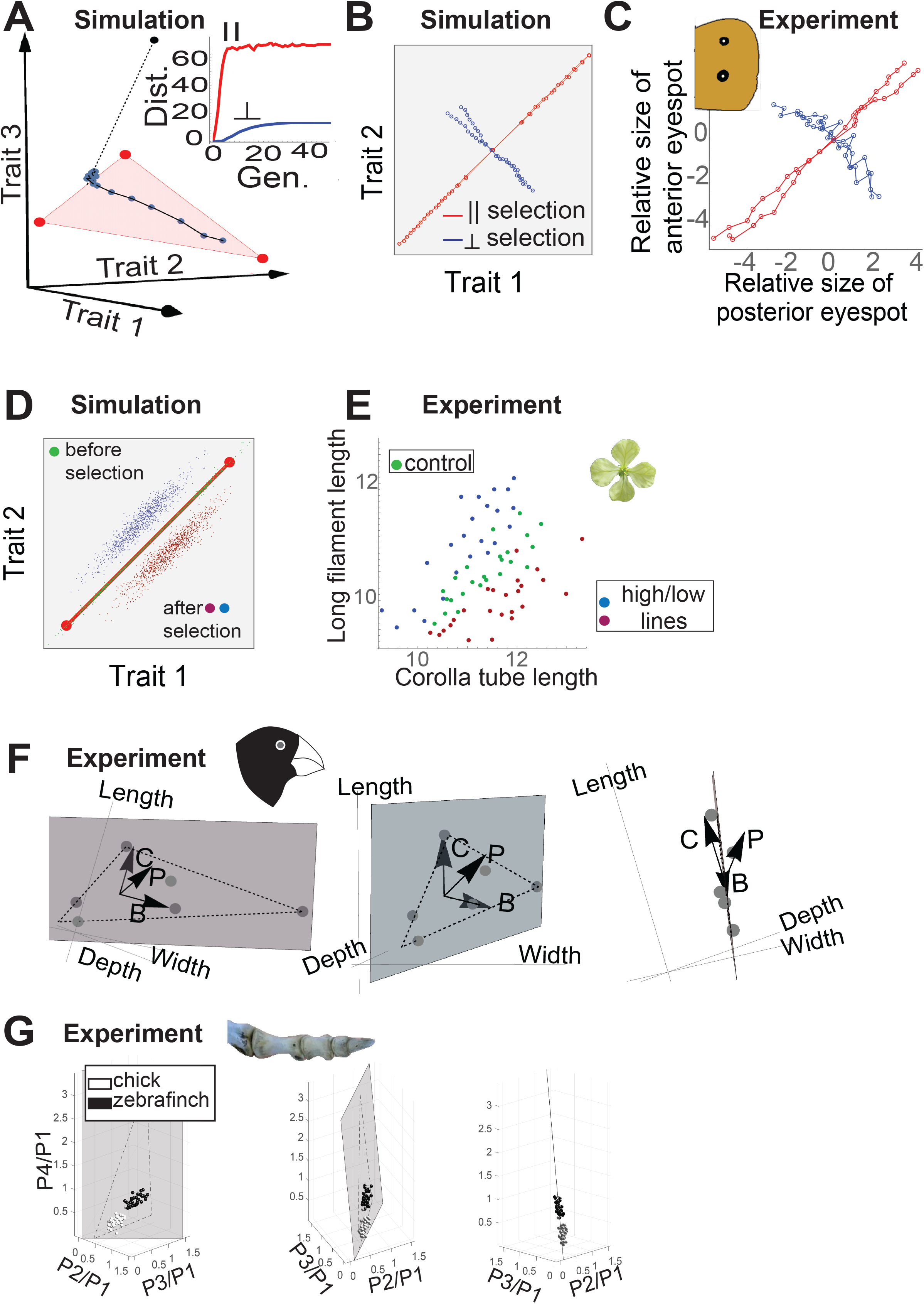
Simulations on artificial selection and population variation agree with a range of experimental results. **(A)** Response to selection towards a target off the front (black point) of an initial set of 100 phenotypes close to archetype 1 in a case of three tasks. The target is equally distant parallel and perpendicular to the front. Each blue point represents the mean phenotype in the population at time steps of one generation of selection towards the target with p=0.1. The population rapidly evolves to the projection of the target on the plane of the triangle, followed by much slower evolution off the triangle. Inset: Parallel (red) and perpendicular (blue) distance from the initial population mean as a function of generation number. **(B)** Response to selection towards a far target along front (red) and perpendicular to the front (blue). Each point represents the mean phenotype at subsequent generations of selection. Initial population was evolved for 10,000 generations with N=100, *μ*=0.5, k=2, infinite-recombination, before artificial selection for another 10 generations. Artificial selection used p=0.5. **(C)** Artificial selection experiments by Allen et. al [31] on butterfly eyespot-size show that the response along the main axis of phenotypic variation of the natural population is larger than the response to selection perpendicular to this axis. The axes are the normalized sizes of the two dorsal eye-spots, and the mean phenotype of the initial population is at the origin. Adapted from [31]. **(D)** Extended simulations of artificial selection perpendicular to the Pareto front results in a line parallel to the front. Initial population was same as in (B), artificial selection was performed for 100 generations. **(E)** Experiments by Conner on radish artificial selection perpendicular to the axis of natural variation results in mean offspring on a line (red, blue) parallel to the axis of natural variation (green). Adapted from [33]. Radish image is modified from https://commons.wikimedia.org/wiki/File:Wild_Radish_flower_(5360402586).jpg. **(F)** Normalized beak dimensions of Darwin’s ground finch species fall approximately on a plane in the space of beak traits. Perturbations of the beak morphogen pathways, BMP, calmodulin, and the premaxillary-bone modification factors denoted B, C and P in the figure, have phenotypic effects (arrows) approximately aligned with this plane (generated using data from [36]). See Electronic Supplementary Material, Section 16 for details. **(G)** Phalanx proportions of different species of birds fall in a triangle. Populations of chickens and zebra-finches show variation aligned with this triangle. Adapted from [11]. Toe image is modified from [11].

Evolution to a target off the front results in rapid motion to the point on the front that is closest to the target (the projection of the target on the plane of the triangle). This is followed by much slower evolution off of the front that stops when perpendicular genetic variation is depleted (Fig 5A).

Simulations of selection in the case of two tasks (Fig 5B) are qualitatively similar to experiments on *Bicyclus anynana* butterfly eyespot size [31], in which artificial selection parallel to the observed suite of variation lead to stronger evolutionary change than selection perpendicular to the suite of variation (Fig 5C).

Additional simulations show that prolonged selection perpendicular to the original front can generate a population that is spread along a hyperplane parallel to the original front (plane for 3 tasks and a line for 2 tasks) (Fig 5D and Electronic Supplementary Material, Section 15, Fig S17D). A similar effect has been observed in experiments on radish morphology (Fig 5E) [32,33].

### The present predictions are supported by a range of experimental data

To further test the present conclusions requires experiments that perturb molecular pathways and measure the resulting phenotypic changes, in relation to the natural suite of variation. A set of studies that exemplify this approach considered the pathways that shape the beak of Darwin’s finches [34–36].

The normalized beak dimensions of different Darwin’s ground finch species fall approximately on a plane (first two principle components account for 99.6% of the variation, Fig 5F), and within that plane on a triangle corresponding to the tasks of eating large seeds, small seeds and pollen/nectar [9,37]. Perturbations of the major beak morphogenic pathways, BMP, calmodulin, and premaxillary-bone modification factors (TGFβIIr, β-catenin and Dkk3), have phenotypic effects that are pleiotropic in a way that is approximately aligned with this plane [36] (Fig 5F). We hypothesize that polymorphisms that affect the expression of these factors are likely to be prevalent and underlie the variation of the beak morphology along the Pareto front.

One can extend this framework to consider different species in a taxon that have comparable traits and share the same tasks. In this case, a prediction of additive, aligned polymorphism structure is that populations within a species will have variation that is aligned with the front defined by the variation of different species [9,38]. A recent example on bird toe-bone proportions showed that different bird species fall on a triangle. The tasks in this case are grasping, walking and scratching/rapturing (Fig 5G) [11]. Populations of chicken and zebra finch individuals fall in a flattened cloud aligned with the triangular Pareto front defined by the other bird species, as predicted (Fig 5G).

## Discussion

We find that allelic polymorphisms that persist under multi-task selection have phenotypic effects oriented along the Pareto front rather than perpendicular to the front. Similarly, polymorphisms that persist have epistatic effects that tend to keep phenotypes on the front.

We also tested the effects of genetic drift on the polymorphism structure. Small population sizes and low competition, both situations with large drift, lead to weaker alignment with the front.

Polymorphism structure is also affected by linkage, which can cause alleles to be carried together. We therefore studied the effect of linkage and asexual mating on the polymorphism structure. We find in simulations with linkage that polymorphisms still tend to align with the front. However, alignment at a given population size is weaker than in the case of free recombination. Instead, allelic polymorphisms have sizable perpendicular components that cancel each other out. Alignment with the front even occurs in simulations with asexual reproduction (no recombination). This is relevant to micro-organisms, and perhaps to tumors, whose gene expression has been suggested to show tradeoffs between tasks [16].

To demonstrate a mechanistic model of tradeoffs and gene-gene epistasis, we analyzed a molecular mechanism that can give rise to polytope-shaped fronts, based on bacterial sigma-factor regulation. Here, k regulators compete over a limiting factor, (eg RNA polymerase). This competition naturally gives rise to a polytope with k vertices in gene expression space. Each vertex corresponds to the expression if there was only one sigma factor present, allowing specialization in a certain task. This mechanism can also provide phenotypic plasticity, the ability of an organism to change its phenotype in response to changes in the environment. Plasticity occurs in this mechanism when environmental signals affect the levels of the regulators through upstream pathways - just as sigma factor activities in *E. coli* are affected by stresses and nutrients. The resulting plasticity moves the phenotypes along the Pareto front and not off of it, reaching optimal solutions in different environments. Thus, in the present picture, plasticity can arise from the same mechanism that provides evolution/selection to a polytope-like front.

The Pareto front in empirical examples from morphology has straight edges when using traits that are traditionally used by morphologists such as bone and tooth areas. However, if these traits are nonlinearly transformed, e.g. from *T_i_* to 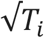, the fronts change from straight to curved. Thus, plotting the data with traits of bone volume or length instead of area would lead to curved fronts. Curved fronts are more difficult for polymorphisms to align to than straight fronts. This raises an interesting hypothesis: the effects of common mutant alleles should be additive in the traits that provide straight fronts (e.g. additive in effects on bone/tooth area rather than volume or length). In this way, additive effects can best keep offspring along the front. In the case of gene expression, in which straight fronts have been observed in log-transformed data [16,17], this prediction means that mutation effects should be additive in log space (multiplicative effects). Indeed, mutation effects often combine in a multiplicative fashion [39,40].

Other ways that phenotypes can evolve to be on a line have been suggested. These mechanisms do not involve multiple tasks, but instead involve migration [41] or consider a fitness landscape with a pronounced ridge [5,6]. These studies do not directly apply to the present case which employs selection in multiple locales with different conditions. More generally, rather than assuming the existence of a fitness ridge a-priori, the Pareto picture suggests a natural explanation for geometries such as lines, triangles and higher-order polytopes in trait space based on the positions of the archetypes that emerge from evolutionary tradeoff between the tasks.

The alignment of polymorphisms with the Pareto front has several implications. First, the progeny of any two parents in the population is likely to be on the Pareto front, as long as the mutations are additive or at least have epistatic effects parallel to the front, such as the epistatic effects described here. Without polymorphism alignment, mating would often result in phenotypes off of the front, which would be at a disadvantage because there exists a potential phenotype on the front which is better at all tasks.

A second feature of the alignment of polymorphisms with the Pareto front is rapid response to selective pressures along the front. Evolutionary response is accelerated towards new selection pressures provided that they relate to the same underlying tasks - and therefore align with the Pareto front. Evidence for rapid response to selection pressures along the main axis of phenotypic variation has been reviewed [42] (Fig 5C).

To further test the present conclusions requires data on many phenotypic traits together with genomic information for a large number of individuals. Such datasets are expected to become more prevalent in the near future [43,44]. It will be fascinating to explore to what extent the effects of common polymorphisms align with the Pareto front of phenotypic traits.

## Author Contribution

HS designed the study, developed and performed the simulations, analyzed the data, and wrote the manuscript. UA designed the study, analyzed the data, and wrote the manuscript. PS participated in developing the simulations. AM participated in analyzing the data. GS participated in analyzing the data and designing the study.

## Acknowledgements

We thank Alon Keinan for fruitful discussions, and the members of the Alon lab for comments on the manuscript. This work was supported by the Israel Science Foundation (1349/15) and the Minerva Foundation. UA is the incumbent of the Abisch-Frenkel Professorial Chair. GS’s work was funded by NIH grant GM115889.

